# High GC Content Causes De Novo Created Proteins to be Intrinsically Disordered

**DOI:** 10.1101/070003

**Authors:** Walter Basile, Oxana Sachenkova, Sara Light, Arne Elofsson

## Abstract

*De novo* creation of protein coding genes involves formation of short ORFs from noncoding regions; some of these ORFs might then become fixed in the population. *De novo* created proteins need to, at the bare minimum, not cause serious harm to the organism, meaning that they should for instance not cause aggregation. Therefore, although the creation of the short ORFs could be truly random, but the fixation should be of subject to some selective pressure. The selective forces acting on *de novo* created proteins have been elusive and contradictory results have been reported. In *Drosophila* they are more disordered, i.e. are enriched in polar residues, than ancient proteins, while the opposite trend is present in yeast. To the best of our knowledge no valid explanation for this difference has been proposed.

To solve this riddle we studied structural properties and age of all proteins in 187 eukaryotic species. We find that, on average, there are small differences between proteins of different ages, with the exception that younger proteins are shorter. However, when we take the GC content into account we find that this can explain the opposite trends observed in yeast (low GC) and drosophila (high GC). GC content is correlated with codons coding for disorder-promoting amino acids, and inversely correlated with transmembrane, helix and sheet promoting residues. We find that for the youngest proteins, i.e. the ones that are most likely to be *de novo* created, there exists a strong correlation with GC and structural properties. In contrast, this strong relationship is not seen for ancient proteins. This leads us to propose that structural features are not a strong determining factor for fixation of *de novo* created genes. Instead these proteins resemble random proteins given a particular GC level. The dependency on GC content is then gradually weakened during evolution.

**Author Summary:** We show that the GC content of a genomic area is of great importance for the properties of a protein-coding *de novo* created gene. The GC content affects the frequency of the codons and this affects the probability for each amino acid to be included in a *de novo* created protein. The codons encoding for Ala, Pro and Glu contain 80% GC, while codons for Lys, Phe, Asn, Tyr and Ile contain 20% or less. Pro and Gly are disorder-promoting, while Phe, Tyr and Ile are order-promoting. Therefore random protein sequences at a high GC will be more disordered than the ones created at a low GC. The structural properties of the youngest (orphan) proteins match to a large degree the properties of random proteins when the GC content is taken into account. In contrast structural properties of ancient proteins only show a weak correlation with GC content. This suggests that even after fixation of *de novo* created proteins largely resemble random proteins given a certain GC content. Thereafter, during evolution the correlation between structural properties and GC weakens.

## Introduction

Proteins without any detectable homologs outside one genome are often referred to as orphans. Orphan protein coding genes can be created by gene duplication, lateral transfer of genetic material and *de novo* gene creation, that are of particular interest, as they are the only source of completely novel protein coding material and present a rare chance for full-frontal functional novelty. Further, studies of the properties of the genes might provide unique insights into the fundamental processes in the formation and selective pressure of all genes since clearly, in the strict sense, all protein superfamilies were once created by a *de novo* mechanism.

Before the genomic era, the scientific consensus held that *de novo* creation of new genes was rare - instead it was believed that the vast majority of all genes were originally generated in an ancient “big bang". However, when the first complete genomic sequences were initially published, this hypothesis was not supported [1]. In fact, to this day, when analysing complete genomes from closely related genomes, a surprisingly high number of orphan proteins persist [2–4]. It has later been shown that some of the initially assigned orphan proteins are not *de novo* created but rather a result of limited phylogenetic coverage of the genomes [5].

Today, supported by the vast amount of complete genome sequences available and improved search methods [6], many of the orphan proteins detected, at least in yeast, appear to be created through *de novo* formation [7, 8]. Some studies indicate that, in yeast, there is a large set of proto-genes: ORFs that remain on the verge of becoming fixed as *bona fide* protein-coding genes in the population [7]. This gives a possible background in explaining how novel proteins can be generated from non-coding genetic material. In other species the genomic coverage has been more limited and therefore the studies have been less detailed.

It is clear that not all identified orphan proteins are *de novo* created. Several reasons for this exists. Some orphans might be classified as such primarily because the relationship with other proteins are missed. This problems is enhanced with a limited amount of closely related genomes and for fast evolving proteins. In addition gene duplication, lateral transfer, gene losses and domain rearrangements also make it difficult to detect the true relationship between all proteins. To accurately detect *de novo* created genes, the availability of several completely sequenced genomes not only from closely related species, but also from a set of numerous and evenly spaced taxa is essential. Even when this is present the best that can be obtained is a set of orphans strongly enriched in *de novo* created proteins.

The availability of complete genomes separated at different evolutionary distances also enables studies at different ages [3,5,7]. Here, a gene can be unique to a specific species, or even to a strain; alternatively it can be present pervasively across a taxonomic group. Even more ancient orphans may be defined as superfamilies that are unique to a kingdom of life [9, 10]. Using methods such as ProteinHistorian it is possible to assign an age to each protein [11].

After *de novo* creation the gene needs to become fixed in the population. The selective forces governing this process have been studied by examining the properties of the *de novo* created proteins that are fixed in the population. Intrinsic disorder, low complexity, subtelomeric location, high *β*-sheet preference as well as other features have been associated with *de novo* created genes and orphan proteins [12, 13]. It has also been proposed that with age proteins (i) accumulate interactions, (ii) become more often essential and (iii) obtain lower *β*-strand content and higher stability [14]. Some aspects of these, such as the fact that orphans on average are short, are likely related to a *de novo* creation mechanism. However, other features, including intrinsic disorder [4, 15], are not obviously related to the *bona fide* gene genesis and could instead be the result of the selective pressure acting during fixation.

In yeast, we have earlier reported that the most recent orphans, i.e. the ones unique to *S. cerevisiae*, are less disordered than the average yeast proteins [3]. Studies enabled by the sequencing of *Drosophila pseudoobscura* provide the opposite picture, i.e. the youngest proteins are more disordered than ancient [4].

To the best of our knowledge the origin of this difference has not been explained. Could the selective forces for *de novo* creation be that disparate between two different eukaryotes, or could the *de novo* genetic mechanisms be different, or is it an artefact caused by evolutionary rates or evolutionary distances between the related genomes? Alternatively, there might exist some genomic feature that is different between drosophila and yeast that could explain the difference of the intrinsic disorder in their orphan proteins. In addition to hugely different sizes and different gene structures, the GC content differs significantly between the genomes of different taxa. The GC content of *Saccharomyces* genomes is roughly 40%, while in *Drosophila* the GC content is 55%.

To obtain a better understanding of the structural properties affecting the *de novo* creation of proteins, we studied the age of proteins in 187 eukaryotic genomes. Significantly more than used in earlier studies. Due to the frequency of lateral transfer in prokaryotic mechanisms age estimates of prokaryotic genes is more troublesome than for eukaryotic genes. Therefore, we focus on eukaryotic organisms in this study.

We find that the most striking difference between young and old proteins is their difference in length. Surprisingly all other properties show a large overlap between ancient and orphan proteins. However, we find that structural features in orphan proteins differ significantly between low-GC and high-GC genes. Orphans in low GC genes are more disordered and have less secondary structure than in high-GC genes. In older proteins this relationship is much weaker, supporting a model where *de novo* creation starts from random non-coding ORFs and then gradually adapts the features of ancient proteins.

## Materials and Methods

### Datasets

To start, protein data for 400 eukaryotic species were obtained from OrthoDB, release 8 [16]. These species are divided into 173 Metazoans and 227 Fungi, for a total of 4,562,743 protein sequences. For each species, a complete proteome was also downloaded from UniProt Knowledge Base [17].

### Age estimate

The ProteinHistorian software pipeline [11] is aimed at annotating proteins with phylogenetic ages. This method requires a phylogenetic tree relating a set of species, and a protein family file, containing the orthology relationships between the proteins of the species in the tree. The pipeline will then assign each protein to an age group, depending on the species tree and the ancestral family reconstruction algorithm used to identify protein families. For our application, we used ProteinHistorian with default parameters, the NCBI phylogenetic tree [18], and protein orthology data obtained from OrthoDB. The OrthoDB method is based on all-against-all protein sequence comparisons using the Smith-Waterman algorithm and requiring a sequence alignment overlap of at least 30 amino acids across all members of an orthologous group. Therefore, the age group can be thought of as the level in the species tree on which a shared sequence of at least 30 AAs first appeared, i.e. it assigns multi-domain proteins to the age of its oldest domains.

One problem that exists using the NCBI phylogenetic tree is the presence of many polytomic branches, especially at the genus level. The cases when more than one of species were present in a multi-furcated branch are problematic, because ProteinHistorian can not distinguish between its proteins being specific to that species and proteins shared among the entire group. To solve this, we converted the NCBI tree to a fully binary by forcing no polytomy on the terminal branches.

### Identification and definition of orphans

Proteins present in OrthoDB are only those with orthologs in at least one other species, i.e. proteins without orthologs (singletons) are not present in OrthoDB. Therefore, to obtain a set of candidate orphan proteins, the complete proteomes of all species were downloaded from Uniprot. Thereafter, BLAST was used to extract proteins not present in the OrthoDB dataset, obtaining 356,884 candidate orphan proteins. However, a large fraction of these proteins are not orphans but are missing from OrthoDB for other reasons, including that they were not present when the database was created or that they have undergone large domain rearrangements. We would assume that truly *de novo* created orphans do not contain domains found in other proteins. Therefore to ensure that we have a unique set of orphan proteins we filtered out proteins with hits in the Pfam-A database, by using hmmscan. We believe that, due to the very stringent criteria used here, the majority of this remaining set is constituted of *de novo* created proteins, and we refer to them as orphans throughout the rest of this paper. These proteins are specific to the species taxonomic level, i.e. we expect not to find them in other species in the dataset, even in the same genus. For *Saccharomyces cerevisiae*, that has several strains in the dataset, we also included the strain specific proteins in the orphan group.

Among the OrthoDB proteins, we defined genus Orphans those that were assigned age = 1 (2 in the case of *S. cerevisiae*, because several strains are present in the dataset) by ProteinHistorian. These proteins are specific to the taxonomic level immediately superior to the one of orphans, i.e. genus Orphans are genus-specific. By this definition, taxonomies represented by a single genus in the dataset have no genus Orphans; for this reason, we selected for our final dataset only those species that have at least one other species within the same genus.

Proteins having the maximum age according to ProteinHistorian were defined as ancient: these proteins are thought to be present in the common ancestor of all Fungi (taxon id = 4751) or all Metazoa (taxon id = 33208). Finally, proteins whose calculated age is between genus orphans and ancient were defined as intermediate.

This final dataset amounts to 1,782,675 proteins distributed across 187 species. On average, 73 orphans were found in a genome, 0.8% of all proteins are defined orphans and 0.6% as genus orphans.

This shows that for most genomes we do a very conservative estimate of the number of orphans. When comparing to earlier published sets of orphans in yeast and drosophila our numbers are significantly lower.

For instance, in *Saccharomyces cerevisiae* (reference strain s288c), we identified 16 orphans and 5 genus orphans, out of 6466 total proteins. As a comparison, in our earlier study we have reported 157 species-specific and 125 genus-specific orphans [19] and Vidal en co-workers reported 143 species-specific (ORFs_1_) and 609 genus-specific (ORFs_2–4_) proteins [20]. In a more detailed view, 50-70% of the proteins earlier described as orphans are here classified as Intermediate. Further, the majority of yeast proteins classified as genus-specific orphans are equally divided between intermediate and ancient. This shows that the identification of exact what proteins are *de novo* created remains a difficult proteins and depends on the genomes included in the study.

Following the same trend, in *Drosophila pseudoobscura* we could identify only 6 orphan proteins, in comparison to the much higher numbers (228) reported previously [4]. Four species were found to have more than 5% of orphans: *Ciona intestinalis* (5.8%), *Colletotrichum gloeosporioides* (6.4%, *Botryotinia fuckeliana* (6.5%) and *Apis mellifera* (7.2%).

In conclusion we do believe that the conservative estimate orphans here is suitable for this study as our primarily aim is not to estimate the exact number of orphans but to examine properties of proteins of different ages. In particular we do believe that among orphans as well as among genus orphans there is a significant fraction of *de novo* created proteins.

### Assigning GC content

To assign the GC content of each gene, we downloaded nucleotide coding sequences (CDS) data for each species from the European Nucleotide Archive [21] and mapped each sequence. The mapping was performed using the Uniprot KnowledgeBase mapping data (ftp://ftp.uniprot.org/pub/databases/uniprot/current_release/knowledgebase/idmapping/idmapping In OrthoDB, each protein has a primary, internal identifier, and a secondary identifier that we could use to search the Uniprot mapping file. The corresponding EMBL identifier was used to download the CDS data from ENA (https://www.ebi.ac.uk/ena/). We could map 1,357,518 out of 1,782,675 proteins (~76% of the dataset). The GC content was then calculated for each mapped protein coding gene individually.

Generally, the GC% of a coding region is higher than that of a non-coding region of DNA [22]; therefore, we expect that, for any given species, the GC of coding segments would be higher than the taxonomic GC. To examine this the genome wide GC content of were downloaded, for each species, from NCBI Genome Reports (ftp://ftp.ncbi.nlm.nih.gov/genorries/GENOME_REPORTS/eukaryotes.txt); Indeed for 94% of the species the CDS sequence is higher than the taxonomic GC. Therefore we find it more relevant to define the genomic GC content as the average, for each species, of the GC of its CDS. Anyhow, results computed for predicted structural properties against the GC content of the genome wide DNA (*taxonomic GC*) are shown in Supplementary material see Fig. ??.

### Predicted properties of proteins

Intrinsic disorder content was predicted for all the proteins by using IUPred in its long disorder mode [23]. A single amino acid residue was then labelled as disordered if its intrinsic disorder was > 0.5. The disorder content of a protein is shown as the percentage of its disordered amino acid residues.

We used SCAMPI [24] to predict the percentage of transmembrane residues of each protein. Low-complexity regions were predicted using the software SEG [25]. For each protein, we indicate as SEG the percentage of residues in low-complexity regions.

PSIPRED ( [26]) was used to predict the secondary structure of all the proteins in the dataset. Here, the secondary structure was predicted using only a single sequence and not a profile. This reduces the accuracy but the overall frequencies should be rather accurately predicted. We annotated each protein with the percentage of its residues predicted to be in each type of structure (alpha helix, beta strand, coil).

### Propensity scales

TOP-IDP [27] is a measure of the disorder-promoting vs. avoiding propensity of single amino acids. For each protein, a single propensity was calculated by averaging the TOP-IDP values of all its residues.

We express the hydrophobicity of each protein as the average score of all its residues using the Hessa hydrophobicity scale [28].

For each protein, we computed the propensity of each amino acid to be in one of the four possible secondary structures (helix, sheet, coil, turn) by using the energy function-based propensity scales proposed earlier [29]. The average propensity for each secondary structure was then calculated for each protein.

### Random proteins at different GC contents

To test whether the studied intrinsic properties (disorder, transmembrane, TOP-IDP and hydrophobicity), as well as the frequency of any given amino acid, were solely dependent on GC content, we used a set of 21,000 random ORFs, generated as follows: at each GC content ranging from 20 to 90%, in steps of 1%, a set of 400 ORFs (equally divided into 300, 900, 1,500 and 2,100 bp long) was generated so that its content of GC was fixed. The ORFs were generated by randomly selecting codons among the 61 non-stop codons. The probability to select one codon given a GC content of *GC_freq_* is set accordingly:

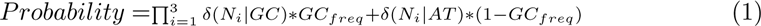

where *N_i_* is the nucleotide of the codon in position *i* and *δ*(*N|GC* is equal to 1 if the nucleotide *N* is guanine or cytosine and zero otherwise, etc. Finally, start and stop codons are added. These ORFs were then translated to polypeptides, and all their intrinsic properties, as well as the frequencies of their amino acid were computed, as described above.

## Results

The assignment of age to all proteins is based on the ProteinHistorian pipeline [11]. In the youngest, orphan group, only proteins that are (a) not present in any other of the 400 eukaryotic genomes in OrthoDB [16] release 8 and (b) that do not share any Pfam-A domain with any other eukaryotic protein are present. Less than 1% of the proteins in the dataset are classified as orphans, see Fig. 1a.

**Figure 1.**
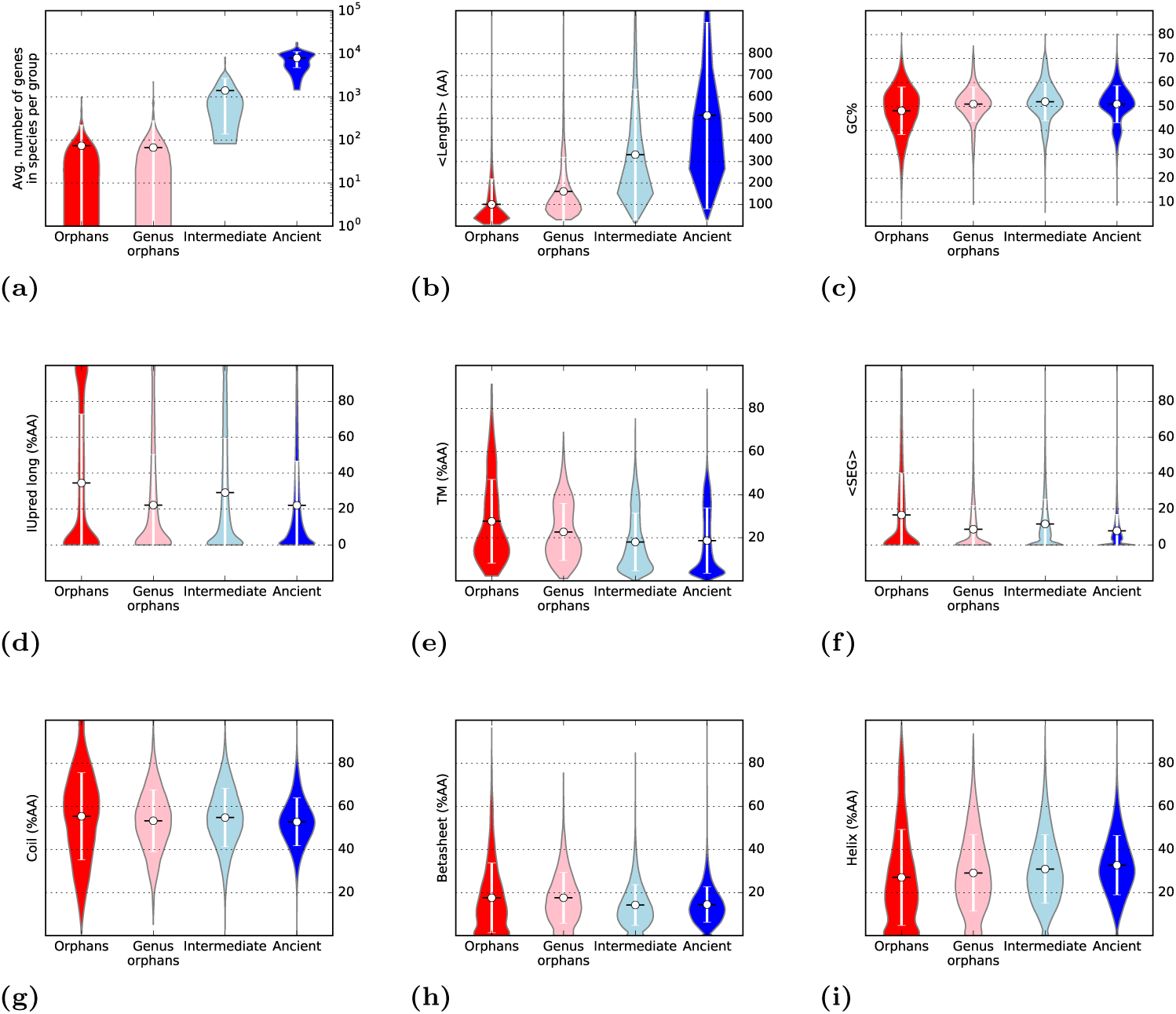
Overview of the proteins assigned to the four age groups in this study. Orphan proteins are proteins unique to one strain/species; genus orphans are found at the immediately superior level (species/genus); Intermediate are found in more general taxonomic levels, but not assigned to be present in the ancestor to all fungi/metazoans. ancient proteins are supposed to be present in the ancestral genomes. In this plots are shown (a) the fraction of proteins belonging to each age group, (b) the average length, in amino acids, (c) the average GC content of the genes, (d) Intrinsic disorder (long) predicted by IUpred (% of disordered residues), (e) percentage of transmembrane residues, (f) fraction of residues in low-complexity regions, (g) fraction of residues predicted to be coil, (h) fraction of residues to predicted to be in a beta sheet and (i) fraction of residues predicted to be in a helix.

In the next group, genus orphans, only proteins that are unique to a genus are included; this group also makes less than 1% of the proteomes. Given that these estimates are significantly more conservative than earlier methods it can be assumed that a large fraction of both orphans and genus orphans.

Finally 10% of the proteins are assigned as intermediate and close to 90% of the genes are ancient. This provides a more strict assignment than most earlier studies see methods for details.

### Orphans are shorter versions of older proteins

The average length of the proteins increases by age, see Fig 1b. The average length is 100 amino acids in orphans, 150 in genus orphans, 300 for intermediate and 500 for the ancient proteins. This highlights the well-established fact that eukaryotic proteins expand during evolution. The expansion can occur by several mechanisms, including domain-fusions [30], additional secondary structure elements [31] and expansion within intrinsically disordered regions [12].

As coding regions on average have higher GC content than non-coding regions [22], it could therefore be expected that GC content would increase by length [32] and therefore by age, but we could not clearly observe this trend, see Fig 1c. It can be however seen how the distribution ofGC in orphans is wider than in ancients, with many genes having less than 40% GC, most likely a consequence of fewer and shorter genes.

Next, we compared predicted structural properties of all proteins see Fig. 1d-i. First it can be noted that none of these properties present a trend as strong as in length. The amount of predicted disorder ranges between 20% and 40% of the amino acids, with the highest average in the orphan and intermediate groups. In orphans, the distribution is bimodal with many completely disordered proteins. Partly this is what is expected for a set of shorter proteins, but certainly it could also indicate there is a preference for a subset of orphans to be disordered.

The fraction of transmembrane residues is on average ~30% in orphan proteins, with a decreasing trend towards ancient (20%). Here, in particular, there are very few young proteins with no predicted transmembrane regions, while these are frequent among the ancient proteins. A similar decreasing trend can be found for low complexity: here orphans have on average 20% of residues in low complexity regions, while less than 10% in ancient proteins. The other structural properties appear to be unaffected by age but with a wider distribution among the younger (and shorter) proteins.

Although some general trends differencing orphans and ancient proteins can be observed, with the exception of length, the relationship actually differs largely between different organisms. For instance when studying intrinsic disorder the orphans and genus orphans of *S. cerevisiae s288c* are remarkably non-disordered (~3% of the amino acids) as shown before [19] see Fig. 2A. The closely related species *Candida albicans* shows a similar trend; see Fig. 2B, while some other Saccharomycetaceae do not.

**Figure 2.**
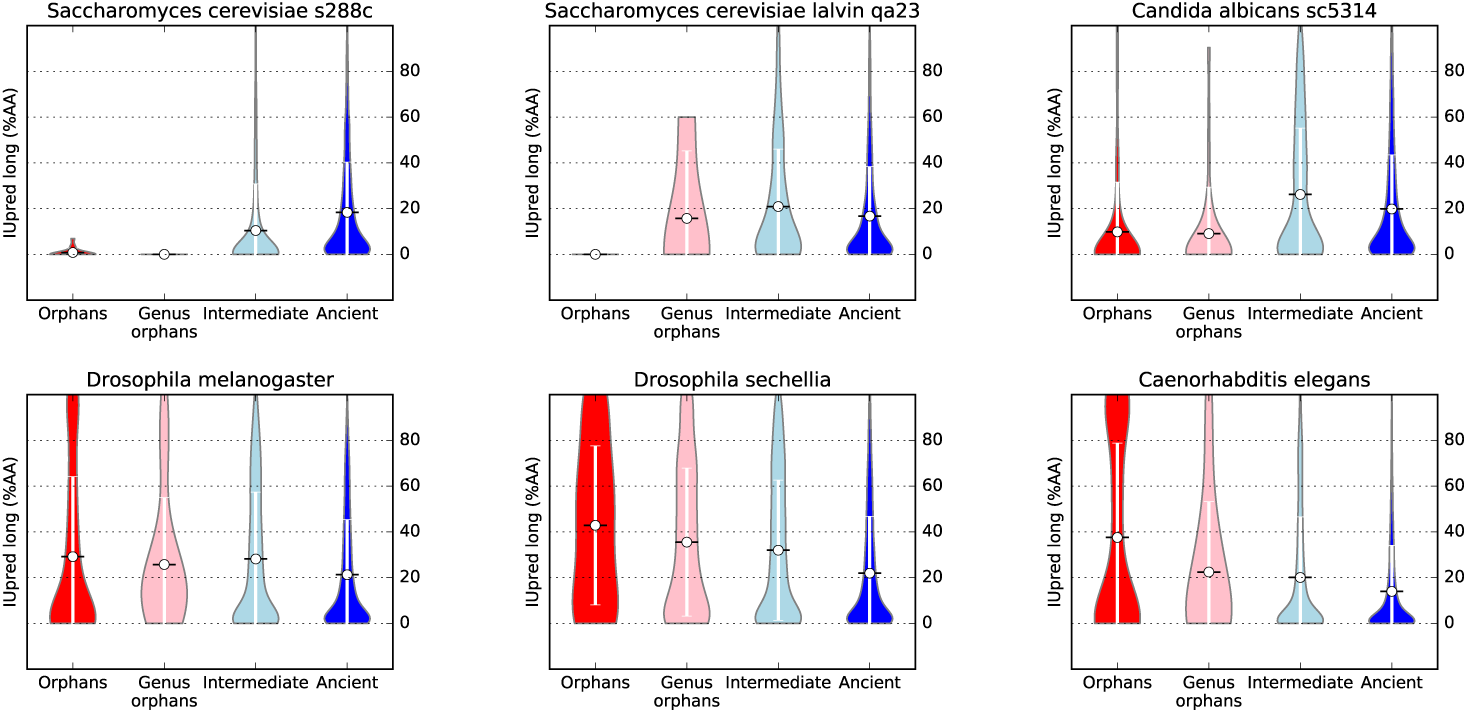
For six selected species (two strains of *S. cerevisiae, C. Albicans, D. melanogaster, D. sechellia* and *C. elegans*), intrinsic disorder (% of amino acid predicted as disordered by IUpred long) is shown as violin plots for proteins in the different age groups.

In contrast, but also consistent with earlier studies [15], *Drosophila* orphans are more disordered than their ancient proteins. orphans and genus orphans in most *Drosophila* genome are more disordered than the ancient, see Fig. 2C. In worm, orphan proteins appear to be consistently more disordered than progressively older ones, across all the considered *Caenorhabditis* species, see Fig. 2D.

In general, it is apparent that the variation among the species is quite large, as in some organism orphans are more disordered than ancient proteins, while in others the opposite appears to be the case. What could possibly explain this difference?

One possibility is that the more complex regulations in animals require more disordered residues in comparison with yeast. But the average disorder content is similar in all eukaryotic species, contradicting this idea. We also noted that yeast is also one of the genomes with lowest GC content (~40%). Therefore, we decided to examine the properties of proteins from different age groups in respect with to their GC content.

### Orphans are more disordered in high-GC genomes

To identify the origin of the different properties of orphan vs. ancient proteins in different organisms, we studied the distribution of structural properties for all genomes against the corresponding GC content see Fig. 3.

**Figure 3.**
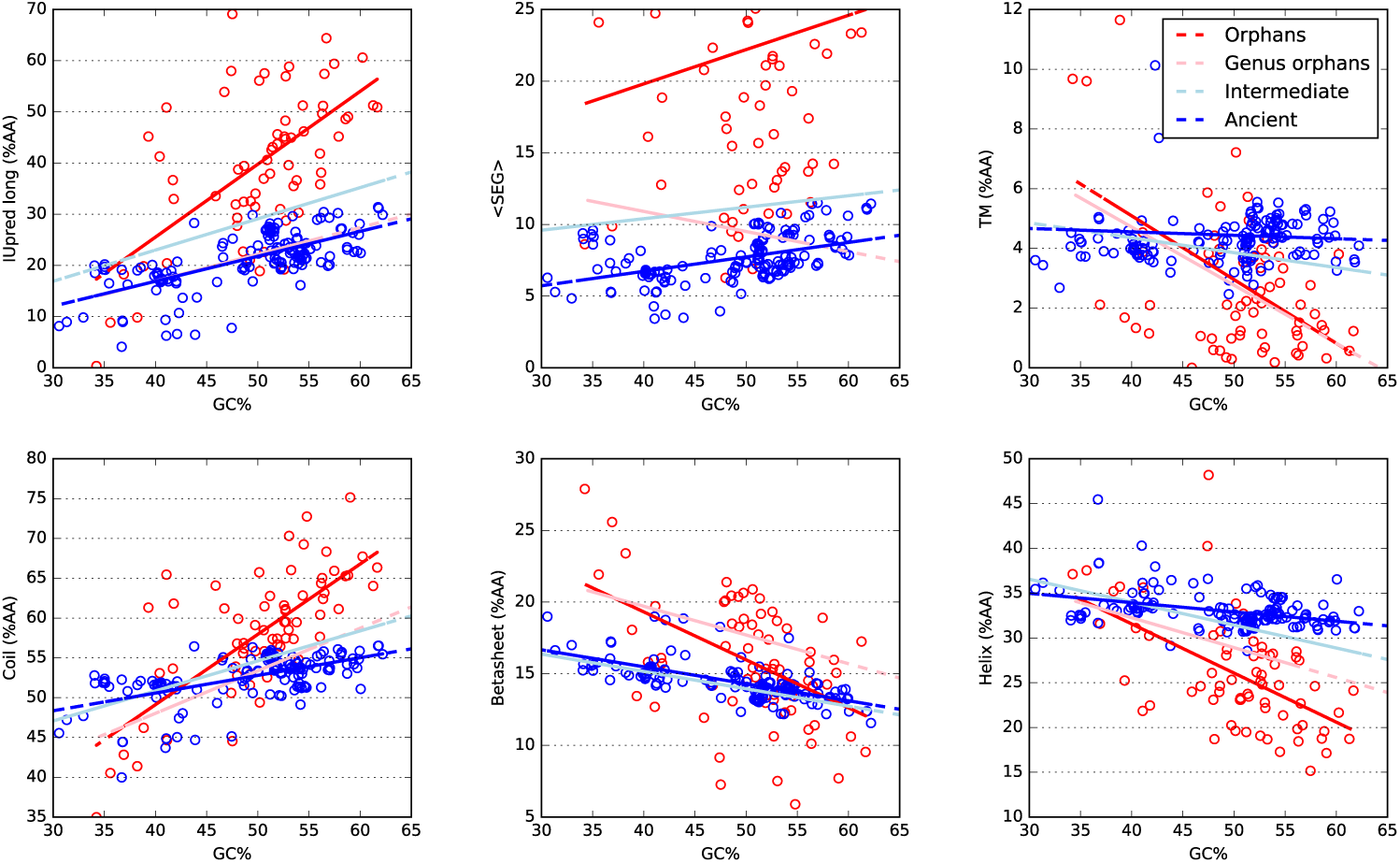
Structural properties of proteins of different ages plotted against the GC content of the genome (coding regions). For clarity only the ancient (blue) and orphan
(red) proteins are shown individually, but the linear fitted lines for genus orphans (pink line) and intermediate ones (light blue) are also shown.

For proteins of all ages, disorder, low complexity and coil frequency increase on average with GC, while transmembrane, helix and sheet frequency decrease. Further, the dependency of GC is clearly stronger for younger proteins, indicating that it is related to the creation of the protein and then gradually lost during evolution.

Notable is that intrinsic disorder shows a clear, directly proportional dependency on GC: higher GC corresponds to more disorder. At the extreme (over 60% GC), more than 50% of the residues are predicted to be disordered in orphan proteins, while for ancient proteins the disorder percentage is about 30%. At low GC (below 40%) the disorder percentages is lower and similar in ancient and orphan proteins (15%). Other structural properties show a similar behaviour; for orphans the transmembrane and coil contents are high in low-GC genomes, while sheet and helix contents are high. For the ancient and intermediate proteins there is a much weaker relationship with GC.

The GC is not constant over a genome. In general coding regions have higher GC than non-coding regions [33]. Further, there are also variation in GC between different regions of a genome, so when a non-coding region is turned into a gene the local GC will decide the amino acid content of the protein. Therefore, it might be more relevant to study the GC of each gene individually.

### A strong relationship between GC and structural properties of orphan genes

In Fig. 4 we show the dependency of structural properties on GC content for individual genes. In addition to the variation for protein of the four age groups, we have examined the structural properties for a set of random proteins generated from codons at a given GC frequency, for details see methods. It can be seen that the structural properties of these random genes are clearly GC dependent.

**Figure 4.**
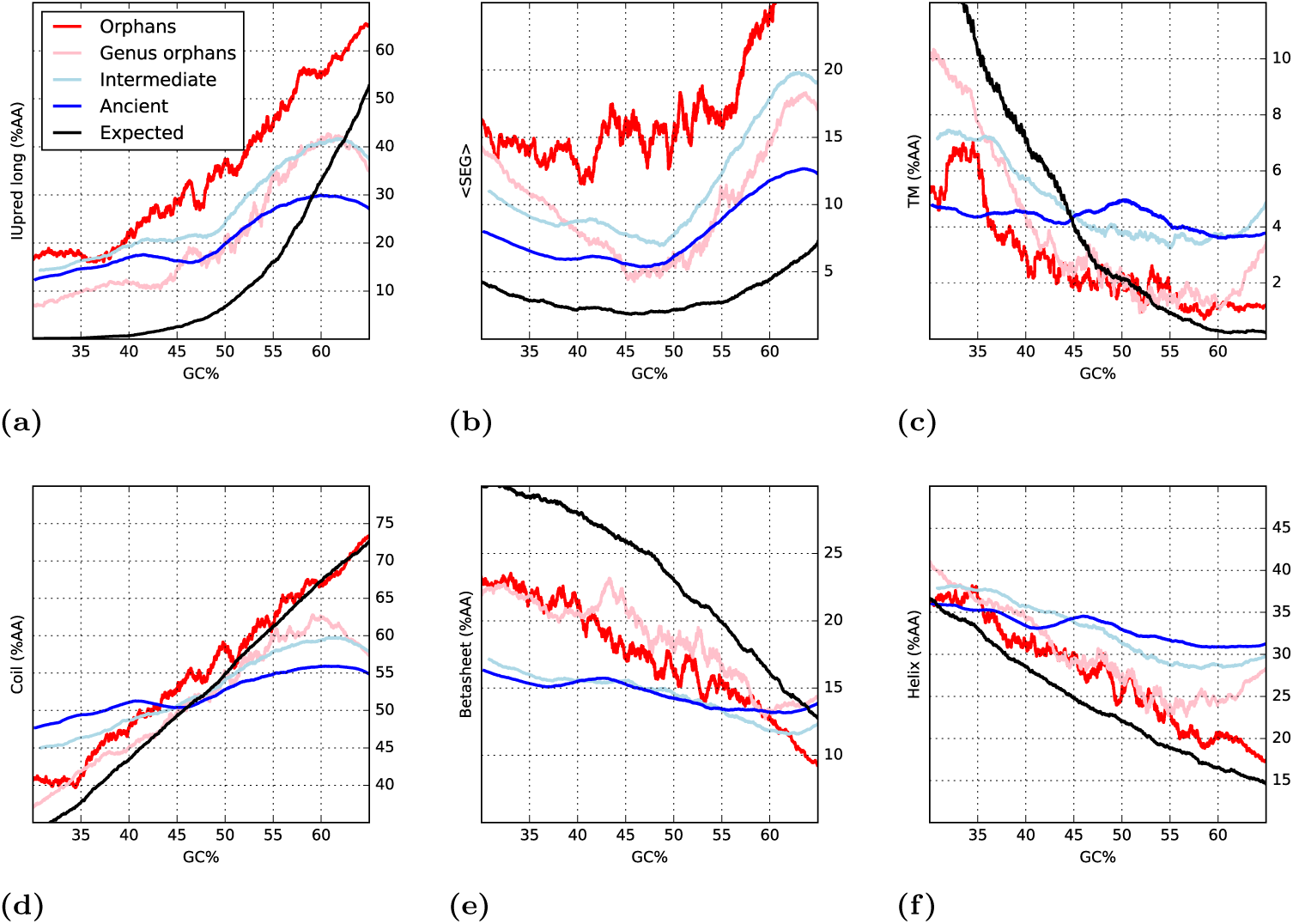
Running averages of predicted structural properties of proteins of different age, orphans (red), genus orphans (pink), intermediate (light blue) and ancient (blue). The black lines represent randomly generated proteins at a given GC frequency.

Orphans, and genus orphans, show a definite dependency of all studied properties on GC, thus indicating that, broadly, orphan proteins appear to be simklar to random protein in their nature, given a certain GC level see Fig. 4. In contrast ancient and intermediate proteins the structural features are only loosely dependent on GC, and they appear to contain less sheet and more helical residues than expected by random.

When studying Fig. 4 in more detail a few notable differences between the random proteins and the orphans can be observed: orphans are more disordered; contain more low complexity regions but fewer sheets independently of the GC level.

It should be recalled that what we describe above is based on *predicted* structural features and they are a reflection of the sequence of a protein. If a certain group of proteins is predicted to be more disordered, or contain more sheets, it is quite likely a consequence of changes in amino acid frequencies, in such a way that the frequency of order-/disorder-promoting amino acids changes.

### Property scales

Next, we studied the relationship of the four age groups of proteins given six different amino acid scales, describing their structural preferences. The difference between the scales and the predicted features used above is that scales are describing general properties and are directly calculated from amino acid sequences, while the predicted features are also based on other properties. For disorder we used the TOP-IDP scale [27], for hydrophobicity we used the biological hydrophobicity scale [28], while sheet, turn, coil and helix propensities were analysed using structure-based conformational preferences scales [29].

In agreement with the predicted values, the average properties in the four groups of proteins are rather similar see Fig. ?? However, when taking the GC content into account all properties of the younger proteins shows a strong correlation with GC, see Fig. 5. To a very large degree the properties of the orphan proteins follow what would be expected from random proteins (black line). However, regardless of GC, orphan proteins are more disordered and hydrophobic, have slightly higher turn and helical propensities, and also lower sheet propensities.

**Figure 5.**
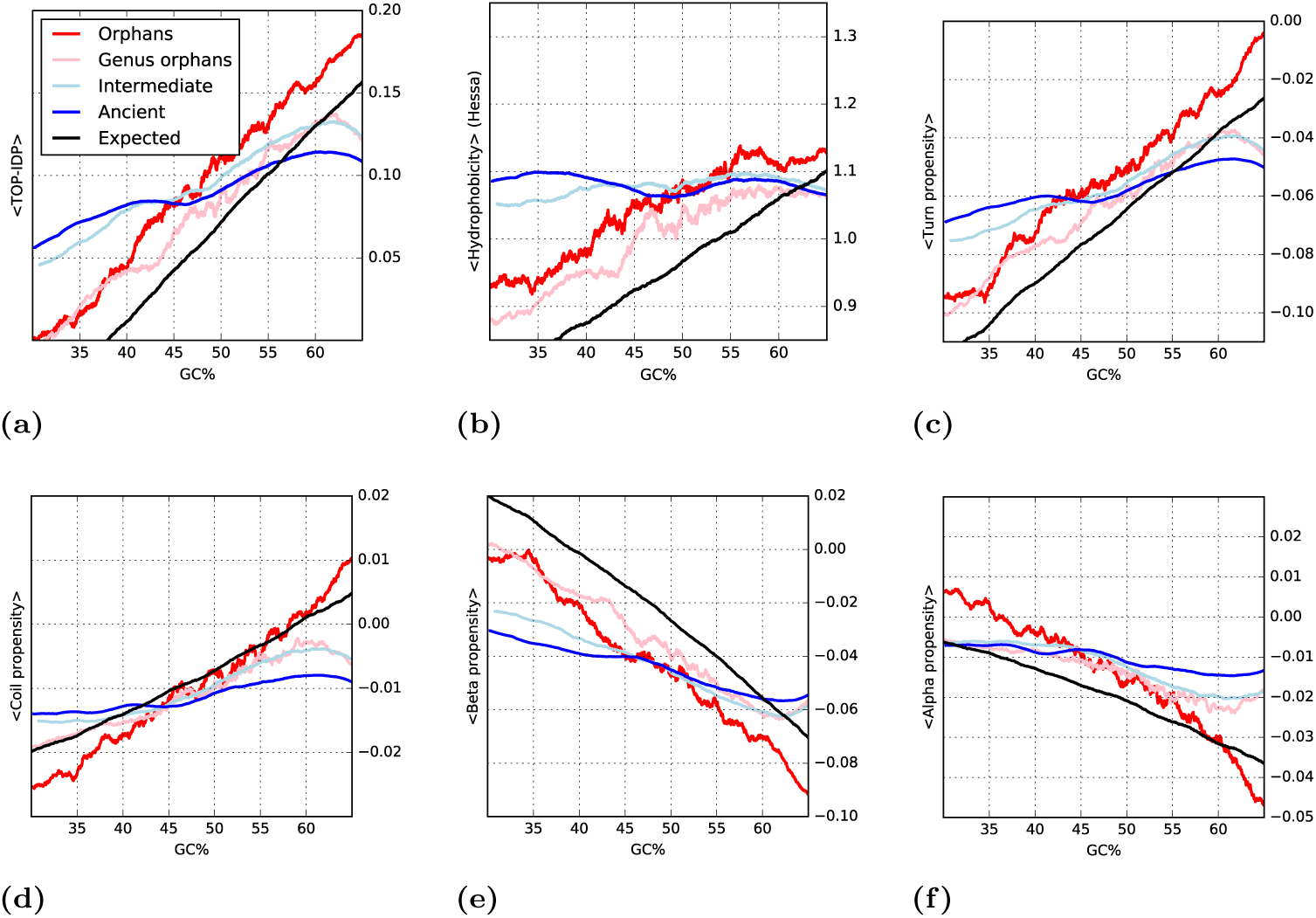
Running averages of structural properties computed from amino acid scales, of proteins of different age, orphans (red), genus orphans (pink), intermediate (light blue) and ancient (blue). The black lines represent randomly generated proteins at a given GC frequency.

Interestingly, also the propensities of the two groups of older proteins change by GC; however, this dependency is much less pronounced than in younger or random proteins. We should remember that amino acid preferences and GC content are coupled both ways: changes of amino acid composition will not only affect the properties but also the codons used and thereby the GC; so it is possible that the relationship between properties and GC for ancient proteins is an indirect consequence of the amino acid preferences and not that the disorder is caused by high GC. The big difference seen between orphan and ancient proteins indicates that, given evolutionary time, the selective pressure to change the GC level is weaker than the selective pressure to change the protein properties.

## Discussion

The GC content affects the codon usage between different genomes [34] and it has been argued that the GC content might be solely responsible for the codon bias [35]. The difference in codon usage causes differences in amino acid frequencies, in such a way that some amino acids are more frequent in higher GC content levels. Obviously, the reverse could also be true, i.e. that high disorder content increases the GC content of a gene. But if this was the case the correlation should be stronger for ancient proteins and not for orphans as we observe here given the fact that ancient proteins should have more time to adjust to the selective pressure. To study the effect of GC content on amino acid frequency we examined the frequency of all 20 amino acids in proteins of different age and GC content.

### The influence of GC on amino acid preferences

How can changes in GC content affect proteins? In a random DNA sequence, the frequency of different codons changes depending on GC, and this, in turns, affects the expected amino acid frequencies. Clearly, the GC content has a strong influence on the structural features of these random proteins (see black lines in Fig. 4 and 5.

In Fig. 6, the expected and observed amino acid frequencies at different GC contents are explored. For most amino acids the observed amino acid frequencies are surprisingly well correlated with what is expected from the codons alone. However, a few notable exceptions exist:

- For Pro, Arg, Trp, Tyr, Phe and Ile, the frequencies in orphan proteins resemble the random proteins and are strongly dependent on GC content, while the frequencies in ancient proteins are much less dependent on GC content. This suggests that there exists a selective pressure to gradually adjust the frequencies of some amino acids to an optimal level.
- Asn and Ala, on the other hand, change in frequency also in ancient proteins, indicating that the selective pressure to change the frequency of these amino acids is lower and it is possible that their frequency is really affected by the GC content of the genome.
- Further, Glu, Gln and Asp are more frequent than expected, at any GC level. Here, the frequency found in orphans is intermediate to what is expected by chance and what is found in ancient proteins. This indicates a gradual adjustment of the frequency of these amino acids during evolution. These amino acids are coded by only two codons, i.e. there exists a selective pressure to increase their frequency to a higher level than the 3.3% expected by chance.
- Finally, Cys and His are less frequent, independently of GC content, in real genes than in random ones, indicating their special roles in protein function and folding as well as their rareness.

**Figure 6.**
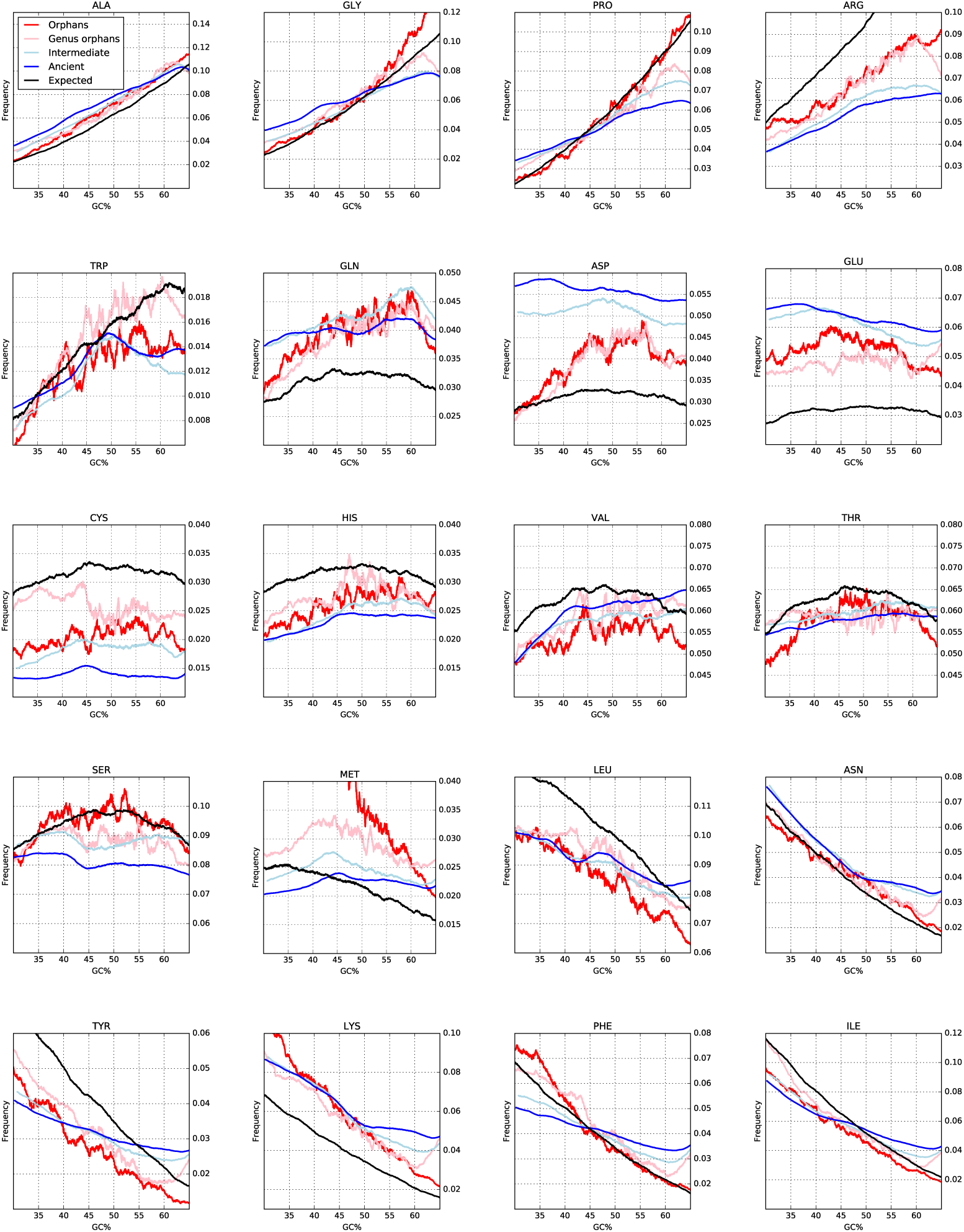
The relationship of each amino acid frequency with the GC content and age of the protein. A black line represents the expected values. The amino acids are sorted by the GC content in their codons.

In Fig. 7 the GC content of the codons of each amino acid is compared with the propensity of that amino acid to be in a certain structural region. Three amino acids, Ala, Gly and Pro are “high GC” amino acids, i.e. they have more than 80% GC in their codons, while five amino acids, Lys, Phe, Asn, Tyr and Ile, have “low GC codons” have less than 20% GC in their codons. The other twelve amino acids show weaker dependency with GC content, see Fig. 7.

**Figure 7.**
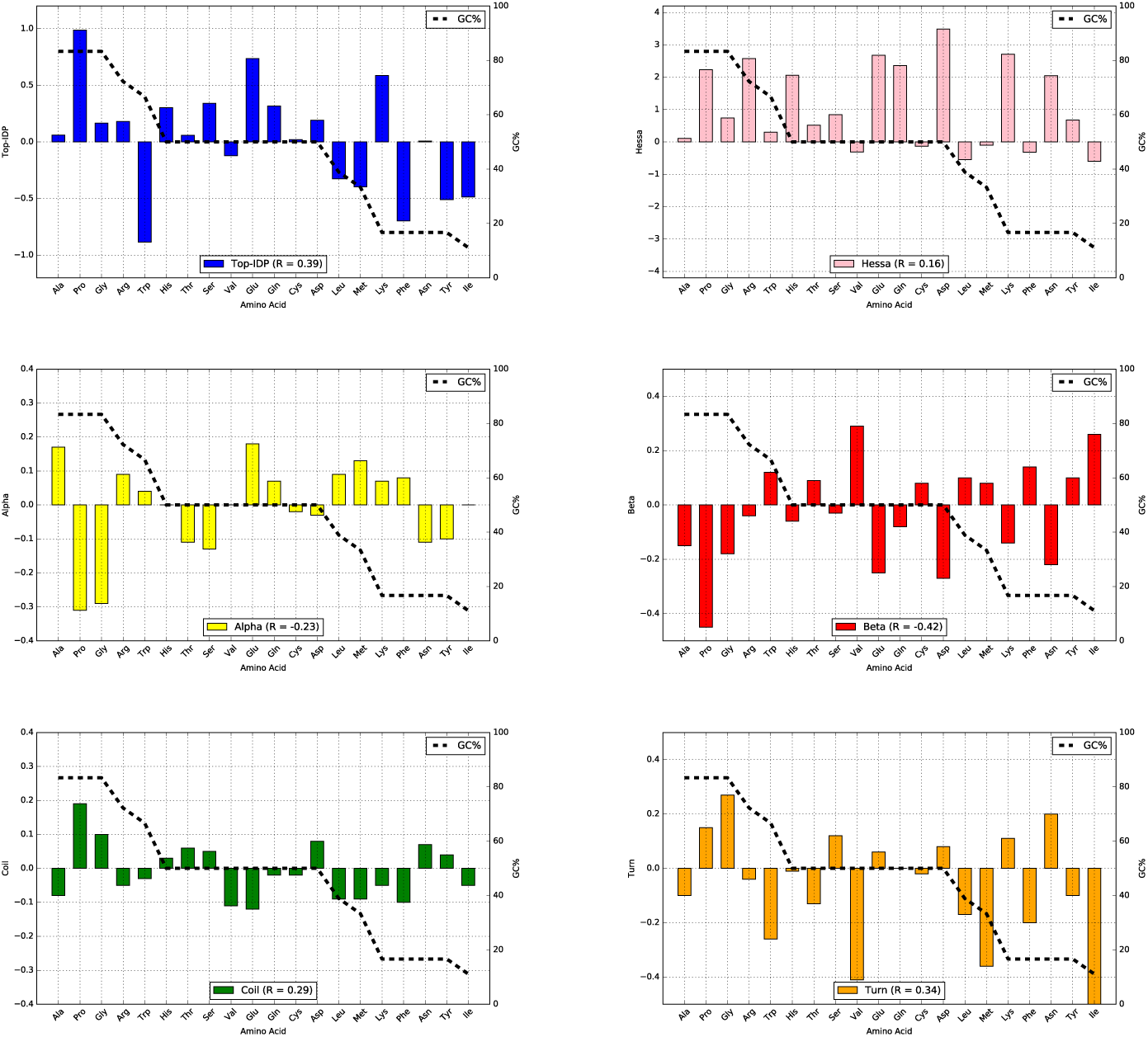
The percentage of GC in all codons encoding an amino acid is plotted as non-filled bars and the values for the different propensity scales as filled bars. (a) TOP-IDP, (b) Hessa transmembrane scale (c-f) Koehl secondary structure preference scale. For each scale the Pearson (R) correlation with GC is also shown.

All three “high GC” amino acids are intrinsic disorder-promoting (high TOP-IDP), while four out of five “low GC” amino acids are order-promoting (low TOP-IDP) residues. Therefore at high GC content, DNA codons coding for hydrophilic, disorder-promoting amino acid are prevalent in any given protein, by simple statistics, while DNA sequences low in GC tend to contain codons for hydrophobic amino acids, associated with low intrinsic disorder.

A comparison between the GC level and structural preferences is shown in Fig. 7. All scales correlate with the GC frequencies with coefficients ranging from −0.42 to 0.39. The strongest correlations are found with *β* propensity (−0.42) and TOP-IDP (0.39) and the weakest with hydrophobicity (0.16). The difference in correlation is mainly caused by the high and low GC amino acids; for example Ile, which is a very strong sheet breaker, is very frequent in turns and rather neutral in most other scales. This contributes to the stronger correlation of disorder and sheet scales with GC compared to other scales. Gly, on the other hand, is a strong helix breaker but not too unfavourable in transmembrane regions, partly explaining why a stronger correlation is observed between GC and TOP-IDP than between GC and hydrophobicity.

so

## Conclusions

We have studied the properties of proteins and their age in a large set of eukaryotic genomes, with a particular focus on the youngest proteins that are most likely to be *de novo* created. As shown before, the youngest proteins are shorter than ancient proteins, but surprisingly we do find that on average for other structural features the young and old proteins are rather similar. We observe that the properties of youngest proteins vary significantly with the GC content. At high GC the youngest proteins become more disordered and contain less secondary structure elements, while at low GC the reverse is observed. We do show that these properties can be explained by changes in amino acid frequencies caused by the different amount of GC in different codons. The influence of this can be seen in the frequency of the amino acids that have a high or low fraction of GCs in their codons, such as Proline.

In a random sequence, the most disorder-promoting amino acid, Pro, only represents less than 5% of the amino acids at 40% GC, but 10% at 60% GC. This actually agrees well with what is observed in the youngest proteins: 5% at 40% GC vs. 9% at 60% GC, see Fig. 6. Interestingly, even ancient proteins show a similar but significantly weaker trend. Here, the fraction of Pro increases from 4.5% to 6%. Similar changes in frequencies can be observed for several amino acids.

On average, young proteins are more disordered than ancient proteins, but this property is strongly related to the GC content. In a low-GC genome the disorder content of an orphan protein is ~30% while in a high-GC genom eit is over 50%, see Fig. 3.

Here we show that GC content of a genome strongly affects the amino acid distribution in *de novo* created proteins. It appears as if *de novo* created proteins that become fixed in the population are very similar to random proteins given a certain GC content. Codons coding for disorder promoting residues are on average richer in GC, explaining the earlier contrasting observations between the low disorder among orphans in a yeast (a low GC organism) and the high disorder among orphans in Drosophila (a high GC organism).

## Acknowledgments

This work was supported by grants from the Swedish Research Council (VR-NT 2012-5046, VR-M 2010-3555) and the Swedish E-science Research Center. Computational resources were provided by SNIC. SL was financed by Bioinformatics Infrastructure for Life Science (BILS).

## Supporting Information

**S1 Fig.**
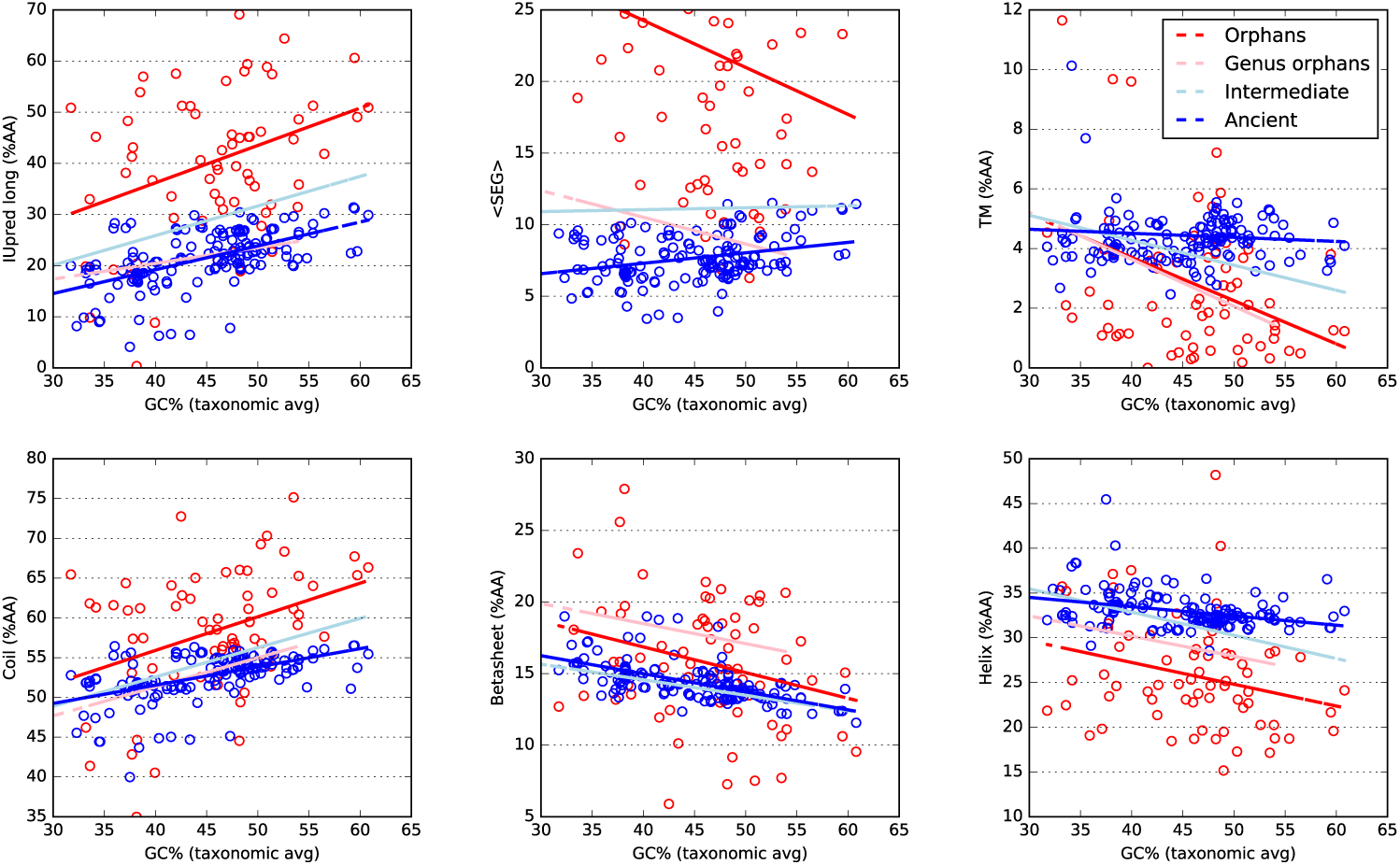
Structural properties of proteins of different ages plotted against the GC content of the genome (entire genome). For clarity only the ancient (blue) and orphan (red) proteins are shown individually, but the linear fitted lines for genus orphans (pink line) and intermediate ones (light blue) are also shown.

**S2 Fig.**
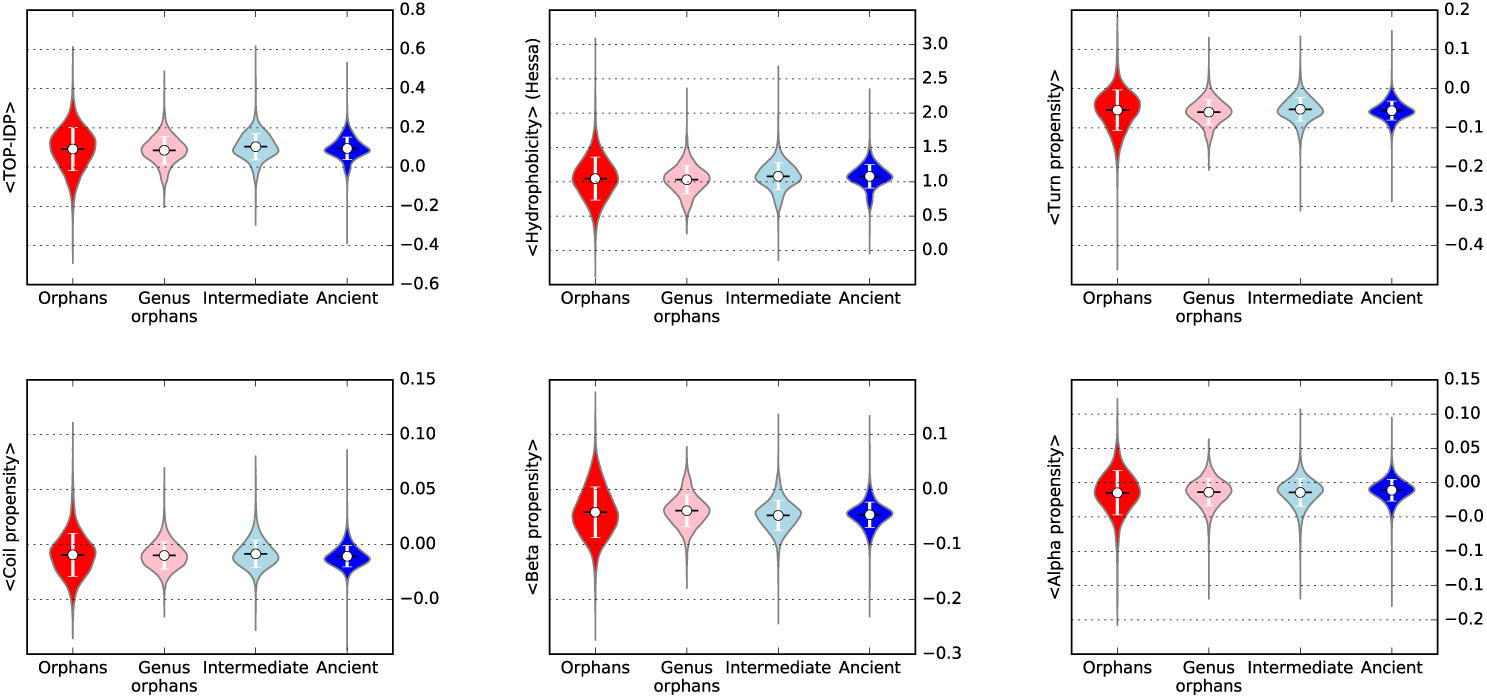
Violin plots showing several properties calculated from propensity scales, as average score. (a) Intrinsic disorder using the TOP-IDP scale, (b) hydrophobicity using the Hessa scale, (c-f) secondary structure preferences.

